# Running ahead of evolution - AI based simulation for predicting future high-risk SARS-CoV-2 variants

**DOI:** 10.1101/2022.11.17.516989

**Authors:** Jie Chen, Zhiwei Nie, Yu Wang, Kai Wang, Fan Xu, Zhiheng Hu, Bing Zheng, Zhennan Wang, Guoli Song, Jingyi Zhang, Jie Fu, Xiansong Huang, Zhongqi Wang, Zhixiang Ren, Qiankun Wang, Daixi Li, Dongqing Wei, Bin Zhou, Chao Yang, Yonghong Tian, Wen Gao

**Author notes:** **Corresponding author:** Wen Gao, Peng Cheng Laboratory, Shenzhen, China., Yonghong Tian, School of Computer Science, Peking University, China., Chao Yang, ICODE and School of Mathematical Sciences, Peking University, China., Bin Zhou, School of Information Science and Engineering, Shandong University, Qingdao, China. These authors contribute equally to this work (Jie Chen, Zhiwei Nie, Yu Wang, Kai Wang, Fan Xu).

## Abstract

The never-ending emergence of SARS-CoV-2 variations of concern (VOCs) has challenged the whole world for pandemic control. In order to develop effective drugs and vaccines, one needs to efficiently simulate SARS-CoV-2 spike receptor binding domain (RBD) mutations and identify high-risk variants. We pretrain a large protein language model with approximately 408 million protein sequences and construct a high-throughput screening for the prediction of binding affinity and antibody escape. As the first work on SARS-CoV-2 RBD mutation simulation, we successfully identify mutations in the RBD regions of 5 VOCs and can screen millions of potential variants in seconds. Our workflow scales to 4096 NPUs with 96.5% scalability and 493.9× speedup in mixed precision computing, while achieving a peak performance of 366.8 PFLOPS (reaching 34.9% theoretical peak) on Pengcheng Cloudbrain-II. Our method paves the way for simulating coronavirus evolution in order to prepare for a future pandemic that will inevitably take place. Our models are released at https://github.com/ZhiweiNiepku/SARS-CoV-2_mutation_simulation to facilitate future related work.

**Justification:** We develop a novel multi-constraint variation prediction framework to simulate SARS-CoV-2 RBD mutations, reaching a peak performance of 366.8 PFLOPS with 96.5% scalability and achieving 493.9× speedup. Our method facilitates the prediction and prioritization of future high-risk variants for the early deployment of drugs and vaccines.

**Performance attributes:** 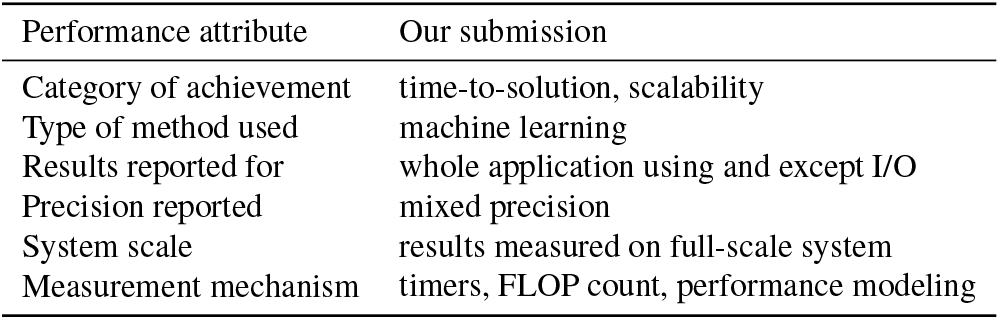

**Overview of the problem:** Coronavirus Disease 2019 (COVID-19) has spread rapidly to more than 200 countries or regions since December 2019. Due to its high infectivity, there have been over 645 million confirmed cases, including approximately 6.6 million deaths, reported by the World Health Organization (WHO) as of December 2022^1^. In addition to being a serious threat to human health, COVID-19 has had a catastrophic impact on the global economy.

---

The virus that causes the pandemic is the severe acute respiratory syndrome coronavirus 2 (SARS-CoV-2) (Figure 1a), which belongs to the genus Betacoronavirus and has nearly 80% sequence similarity with the severe acute respiratory syndrome coronavirus (SARS-CoV) (Lamers and Haagmans 2022; Coronaviridae Study Group of the International Committee on Taxonomy of Viruses 2020; Zhou et al. 2020).

**Figure 1.**
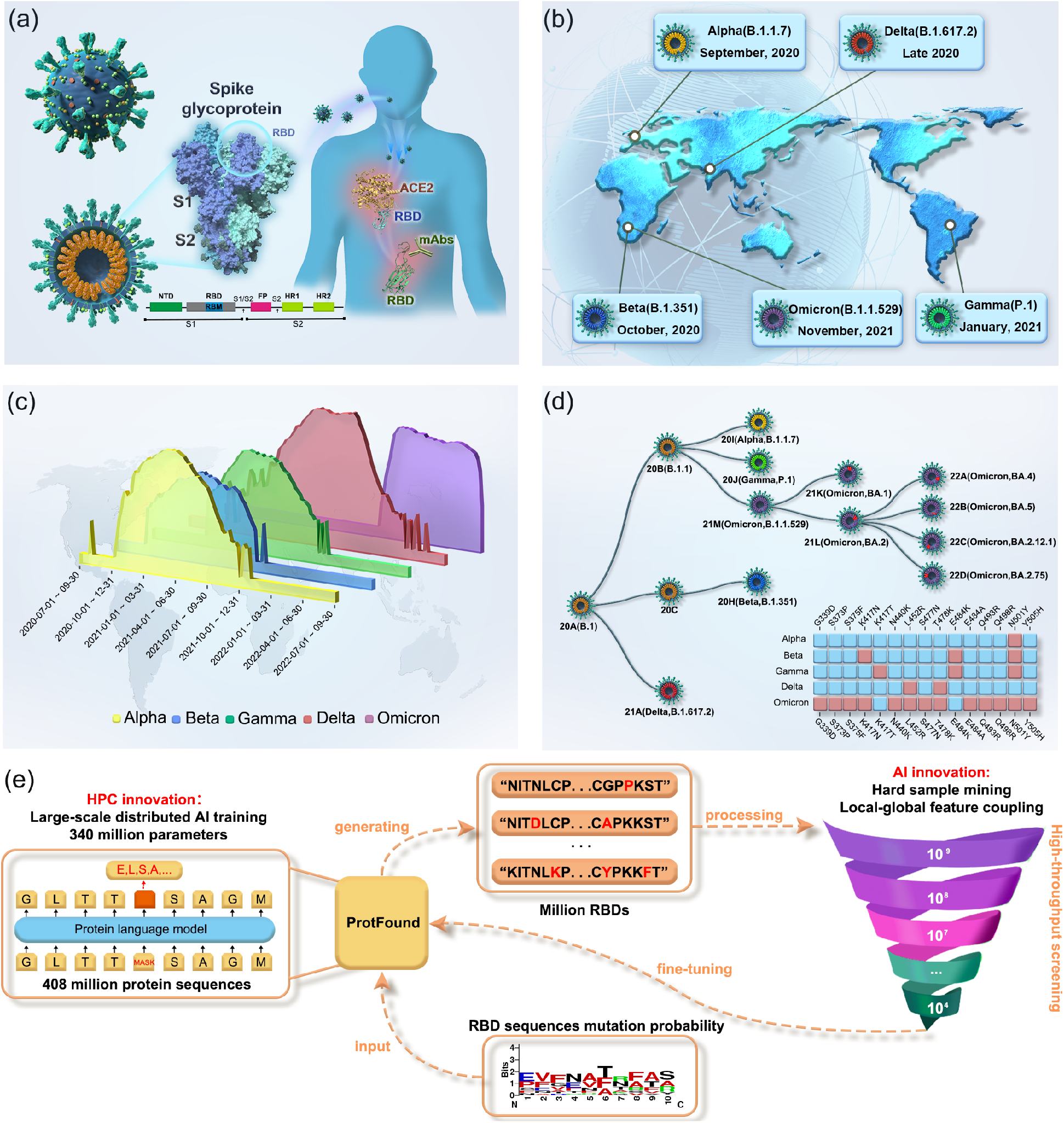
Overview of the problem and our solution. (a) The structural diagram of SARS-CoV-2, in which the RBD on the spike protein is an important region to which hACE2 and the majority of neutralizing antibodies bind. (b) The approximate detection time and places of the five VOCs (Alpha, Beta, Gamma, Delta, and Omicron). (c) Waves of infections caused by the five VOCs from the outbreak of COVID-19 to the present. (d) The phylogenetic tree of SARS-CoV-2 VOCs and the comparison of the variation sites of the five VOCs in the RBD regions. (e) Our methodology for simulating the viral mutation in the RBD. With the support of an HPC optimization strategy that integrates software and hardware, a protein language model (ProtFound) is efficiently pretrained for the generation of RBD mutations. With reference to the mutation frequency of each mutation site in the RBD in the real world, ProtFound can generate billions of RBD variants. These variants are sequentially screened by binding affinity with hACE2, and antibody escape capability. The screened variants are used to fine-tune the ProtFound generator. The fine-tuned ProtFound model is more likely to generate viral variants with higher binding affinity to hACE2 and better capability for antibody escape.

As the pandemic enters its third year, SARS-CoV-2 has been creating waves of infections around the world (Figure 1b,c) (Callaway et al. 2022) due to the high mutation rate of this RNA virus. Which potential SARS-CoV-2 variants may become the next VOCs? Do we need to develop new vaccines to deal with new variants? In what direction will the virus evolve? Shall we just give up as a society and hope that the virus will finally fade away? These are the inconvenient questions that every country on this planet must answer.

Before the current pandemic, the best-known Betacoronaviruses are SARS-CoV and Middle East respiratory syndrome coronavirus (MERS-CoV), which have relatively more severe clinical symptoms than most coronaviruses, which can infect humans but cause only mild symptoms (Yin and Wunderink 2018; Drosten et al. 2003; Zaki et al. 2012; Su et al. 2016; Lu et al. 2020). In the past two decades, the viruses mentioned above have led to two epidemics: SARS (2002) and MERS (2012)(Lu et al. 2020). SARS-CoV-2 can also infect the human respiratory system, but has a much higher infection rate than that of SARS-CoV or MERS-CoV (Walls et al. 2020; Wrapp et al. 2020).

Three sets of proteins, including structural proteins, nonstructural proteins, and accessory proteins, are encoded by SARS-CoV-2 (Lamers and Haagmans 2022) (Figure 1a). There are four main classes of structural proteins, namely, spike protein (S), nucleocapsid protein (N), membrane protein (M), and envelope protein (E), which support the structure of the virus in terms of shape or function (Wu et al. 2020; Lamers and Haagmans 2022). In particular, in addition to their high similarity in sequences, SARS-CoV-2 and SARS-CoV have the same mechanism of infecting host cells, that is, binding to the host entry receptor angiotensin-converting enzyme 2 (hACE2) (Zhou et al. 2020; Wan et al. 2020; Hoffmann et al. 2020; Li et al. 2003). During infection, the trimeric S protein is cleaved by host proteases into the N-terminal S1 subunit and the C-terminal S2 subunit. The receptor-binding domain (RBD) is an important component of the S1 subunit (Figure 1a) that is responsible for binding to hACE2, and is the primary binding target for neutralizing antibodies (NAbs) (Belouzard et al. 2009; Wrapp et al. 2020; Lu et al. 2015; Chi et al. 2020). Therefore, the S protein plays a key role in viral infection and the immune evasion process (Gallagher and Buchmeier 2001; Simmons et al. 2013).

SARS-CoV-2 continues to mutate with a high mutation rate (Duffy 2018) and has evolved into five main variants of concern (VOCs)^2^ as of May 2022: B.1.1.7 (Alpha), B.1.351 (Beta), P.1 (Gamma), B.1.617.2 (Delta) and B.1.1.529 (Omicron) (Figure 1b,c). These SARS-CoV-2 variants with novel spike protein mutations have created waves of infections and reinfections across the globe (Figure 1d). It is vitally important to identify early (Obermeyer et al. 2022) or, even better, to predict dangerous viral mutations that may enhance viral fitness including binding affinity, viral infectivity, or immunity escape.

The Global Initiative on Sharing All Influenza Data (GISAID)^3^ (Shu and McCauley 2017) has recorded more than 14 million SARS-CoV-2 genomes submitted by scientists around the world. This large number of genomic sequences presents an excellent opportunity to study the spread and evolution of SARS-CoV-2. Computational methods such as the Gillespie algorithms can be used to simulate realistic substitution patterns of closely related genomic large-scale datasets, e.g., simulators targeting gene trees, ancestral recombination graphs, or phylogenetic trees (Beiko and Charlebois 2007; Hudson 2002; Laval and Excoffier 2004; Ewing and Hermisson 2010; Rambaut and Grass 1997; Fletcher and Yang 2009; Sipos et al. 2011; De Maio et al. 2022; Shchur et al. 2022). Artificial Intelligence (AI) models can also learn hidden evolution patterns from the huge number of virus sequences submitted, prioritizing future potential viral mutations that could introduce the next VOCs (Chen et al. 2020; Mohamed et al. 2021).

As shown in Figure 1a, the RBD region of the spike protein is an area of concern because it has a high mutation rate, which can significantly affect binding to hACE2, as well as antibodies. In this work, we simulate RBD mutations by learning, generating, screening, and fine-tuning based on pretrained protein language models as shown in Figure 1e. A multi-constrains variation prediction (MCVP) framework is designed to learn from millions of RBD sequences and experimental measurements of binding affinity between single RBD mutations and hACE2/antibodies. MCVP utilizes active learning based on a pretrained protein language model. This high performance computing (HPC) driven work can evaluate RBD mutations based on protein expression, binding affinity, and antibody escape to ultimately provide assistance in the fight against SARS-CoV-2.

## Current state of the art

### Predictive modeling of SARS-CoV-2 variants

During the pandemic, studies have emerged with a variety of focuses and models to predict the mutation of SARS-CoV-2. For example, a renewal-equation-based model was used to describe the adaptive evolution among multiple variants of SARS-CoV-2 including R.1, Alpha, and Delta, and then to predict the dominant variants in Japan before the start of the Tokyo Olympic Games (Ito et al. 2021). Furthermore, some work sought to accurately predict the fitness of SARS-CoV-2 variants, which was used to characterize how efficiently the virus produces infectious progeny. A computational model named SpikePro (Pucci and Rooman 2021) was designed to predict the fitness of SARS-CoV-2 from the sequence and structure of the spike protein in order to allow the identification of new dangerous variants. PyR_0_ (Obermeyer et al. 2022), a hierarchical Bayesian multinomial logistic regression model, was developed to infer relative transmissibility of lineages, forecast future lineage proportions, and identify mutations relevant to fitness. Deep Learning (DL) models have recently been shown to perform well in predicting variant adaptation. Specifically, a three-dimensional convolutional neural network (3D CNN) based on spike dinucleotide composition representation was used to learn the human adaptation of existing coronaviruses and predict the adaptation of SARS-CoV-2 VOCs (Li et al. 2022).

Language models have been used to decipher the genetic sequences of virus. For example, a Transformer-based discriminative model was trained with SARS-CoV-2 genetic sequences to predict potential mutations that may lead to enhanced virus transmissibility (Wu et al. 2021). Language models have also been applied for protein prediction tasks, as common protein motifs and domains can be analogized to words, phrases, and sentences in human language (Ofer et al. 2021; Trifonov 2009; Strait and Dewey 1996; Yu et al. 2019). Motivated by the success of masked language models such as BERT (Devlin et al. 2018), we design a pretrained protein language model for comprehensive variant prediction, aiming to simulate circulating viral mutation and predict potentially risky variants. In this work, we pretrain our protein language model on a large-scale set of protein sequences using a supercomputer with exascale AI training capabilities and further perform fine-tuning and multiconstraint screening on RBD sequences of the spike protein in SARS-CoV-2 to generate possible future variant branches.

### Large-scale language model training

The existing state-of-the-art language models, especially various BERT variations (Devlin et al. 2018; Yang et al. 2019; Howard and Ruder 2018; Liu et al. 2019; Lan et al. 2019) with Transformer as the core, have achieved outstanding performance in many fields. Recently, some works have emerged with a focus on transferring language models to large-scale protein representation learning, e.g., ESM (Rives et al. 2021) and ProtTrans (Elnaggar et al. 2022), which were trained on the Summit supercomputer, and demonstrated that large-scale pretrained language models can capture latent grammar of protein sequences to a certain degree (Elnaggar et al. 2022).

Mini-batch stochastic gradient descent has been found to be very effective for large-scale learning (He et al. 2021). However, updating the parameters in small batches makes the optimization unstable (Li et al. 2020). For large-scale datasets, large-batch training with data parallelism has found increasing popularity (Liu et al. 2019), as it can improve data communication and hardware utilization of a model. However, how to set the best batch size is a complex optimization problem. Some works (Hoffer et al. 2017; Keskar et al. 2016; Goyal et al. 2017; Osawa et al. 2022) have reported that increasing the batch size beyond a certain point can result in poor generalization performance.

### Innovations realized

#### Overview of MCVP

Our proposed multi-constrains variation prediction (MCVP) framework is a heterogeneous system for simulating the effect of the RBD mutations on the fitness of SARS-COV-2 viruses. This system includes 1) a pretrained protein language generative model for RBD mutation generation, 2) an RBD and hACE2 binding affinity prediction model for selecting RBD mutants that have higher binding affinities than the wild type, and 3) an immune escape prediction model for selecting RBD mutants that are more likely to evade antibody attacks.

The training and validation data for the system are collected from various authoritative resources. We download protein sequences from the UniRef database (Suzek et al. 2007) for the training of the protein language model. We download data related to SARS-COV-2 from the GISAID database, which includes more than 14 million genome sequences of SARS-CoV-2 for rapidly sharing. The S protein sequences are obtained from GISAID, then the RBD region sequences are segmented for model fine-tuning and analyzed for the probability of the mutation rate at each position. SARS-COV-2 VOC defining mutations are obtained from https://outbreak.info/.

#### The workflow of MCVP

We design the MCVP framework to follow the workflow as shown in Figure 2a. The first module of MCVP is a Transformer-based language model, hereafter called ProtFound (Protein Foundation Model). ProtFound is trained with the UniRef90 dataset, including approximately 144 million protein sequences. All protein sequences are chopped into lengths of 256, as the RBD region of the spike protein S1 consists of 201 amino acids within the location range of 331-531 (Starr et al. 2020). The structure of ProtFound is similar to that of BERT, but there is no classification token. BERT is a bidirectional model for natural language processing that attempts to reconstruct corrupted tokens. For protein language modeling, 15% of each input protein sequence is masked. During the training process, ProtFound reconstructs the masked amino acids. After training, ProtFound can learn protein embeddings that captured some of the biophysical features of the protein sequences.

**Figure 2.**
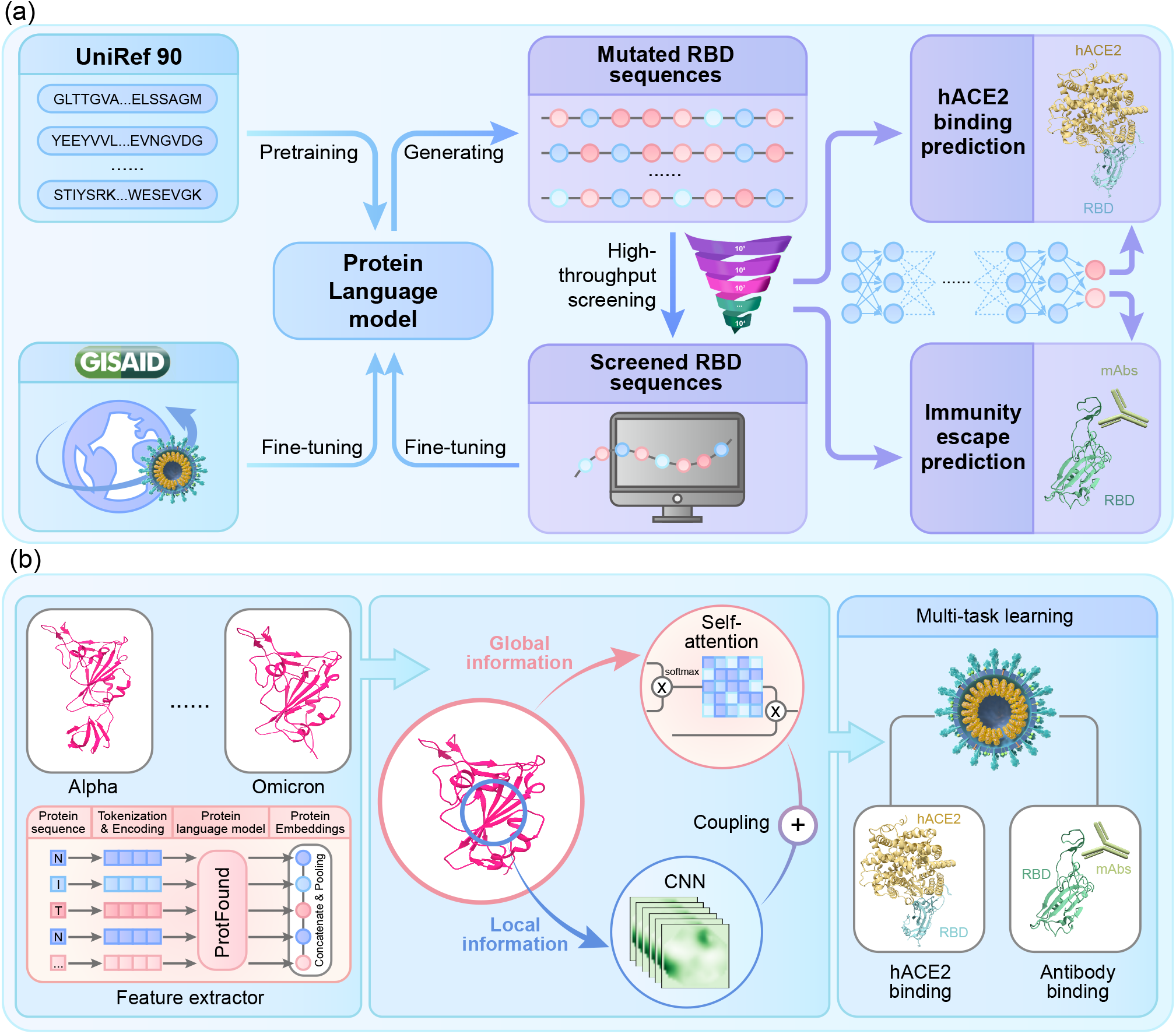
(a) The workflow of our multi-constrains variation prediction (MCVP) framework. It consists of four modules, i.e., pretraining, fine-tuning, generation, and high-throughput screening. (b) Two transfer-learning models for high-throughput screening. Three modules make up the whole processing workflow: a feature extractor module, a feature refinement module, and a downstream task module. The protein embeddings learned by ProtFound are further refined through the coupling of global and local features. Finally, neural networks are trained for two different downstream tasks.

We use ProtFound in two ways. First, we design an RBD-variation-generating module. Specifically, we fine-tune ProtFound with RBD sequences truncated from the spike protein sequences which were downloaded from GISAID. Subsequently, we generate new RBD mutations by generating missing amino acids from a masked RBD sequence selected as the starting sequence. Second, as a protein embedding extractor, ProtFound provides meaningful vector representations of RBD mutations. These embeddings are used as the inputs to a binding affinity prediction model, and an immunity escape prediction model. The above models are essential in selecting RBD mutations that are more advantageous in the sense of virus fitness and survival because of higher binding affinities and immune evasion.

We employ ProtFound to generate millions of RBD mutations with Pengcheng Cloudbrain-II. Subsequently, the two AI filters are used to screen the various generated variants of the RBD based on hACE2 binding affinity and immunity escape respectively in a high-throughput manner. The *in silico* screening is designed to simulate the evolution of SARS-CoV-2 in nature. Therefore, the variants passing this screening could be considered evolutionarily more advantageous. After completing one round of mutation simulation, the selected variants are used as training samples to fine-tune the mutation model ProtFound, which forces the model to learn the characteristics of those variations that are more likely to survive the evolutionary selection. By repeating this procedure, ProtFound is guided to generate variants that are more likely to have evolutionary advantages, thus enabling the simulation of SARS-CoV-2 RBD mutation generation.

As shown in Figure 2b, the protein embedding generation process starts with the tokenization of a protein sequence and the addition of the positional encoding. The resulting vectors pass through ProtFound to create context-aware embeddings for each amino acid, which are the last hidden state of the Transformer’s attention stack. Then these embeddings are concatenated and pooled along the length-dimension to obtain a fixed-size embedding irrespective of the sequence length. In MCVP, the two AI predictors are developed, based on the sequence embeddings extracted by ProtFound. The first is a binding affinity predictor designed for forecasting changes in binding affinity between the mutated RBD and hACE2. The second predictor can be used to evaluate the comprehensive antibody escape capability of the variants through antibody escape prediction.

#### Generation of variants

A variant generation module is designed based on the ProtFound model. Essentially, the ProtFound model has learned the general properties of proteins through self-supervised learning on billions of protein sequences. Then, by fine-tuning ProtFound on millions of RBD sequences, the model is exposed to the subtle amino acid changes in the RBD region of the S1 proteins that are present in the GISAID submissions. We conclude that the final converged model should be able to generate RBD like sequences that would be very likely to new RBD mutations as long as proper constraints are satisfied, e.g., increased binding affinity to hACE2 and increased antibody evasion.

We generate RBD variants by performing the following steps. 1) Spike protein sequences are downloaded from the GISAID database, and the sequences in the RBD region are extracted. 2) Training datasets are created from the data processed in step 1. For each VOC, we create a training dataset using all RBD sequences from the Spike protein sequences that are submitted before the first appearance of that VOC. 3) The ProtFound model is fine-tuned using the training dataset. 4) A variation probability for each position in the RBD is calculated using the training dataset. 5) The variation probability is used to create masks for each position in the RBD. 6) The variant generation module is used to create amino acids at the masked positions.

#### High-throughput screening

Once we have generated a large number of mutation sequences, the next step is to simulate the selection pressure faced by viruses through high-throughput screening. Two screening principles are adopted to perform the progressive filtering of the generated mutations. First, since the main receptor for entering human cells is hACE2, the affinity between the virus RBD and hACE2 is an important indicator for the viral entrance. In other words, future variants should maintain ideal binding affinity with hACE2. Second and more importantly, various studies have shown that VOCs can escape binding to antibodies. Therefore, we design a model to predict binding affinity and a model to predict the immunity escape of the variants. These two models are built with ProtFound as the backbone and are developed based on transfer learning.

#### Simulation of circulating mutations

SAR-CoV-2 is constantly evolving within a host. As a result of evolutionary pressures, viruses tend to mutate to acquire stronger fitness, including better binding affinity, and stronger antibody escape capabilities. We simulate the mutation of SARS-CoV-2 through high-throughput screening and fine-tuning. In each round of stimulation, we use AI models to select those variants that are predicted to retain ideal binding affinity and stronger antibody escape capabilities. The screened variants will then be used for next round of fine-tuning of ProtFound. These steps complete the *in silico* mutational simulation of SARS-CoV-2 RBD.

#### HPC strategy design

For large-scale distributed AI training, the main goals are to optimize the throughput and speed up network convergence. Pengcheng Cloudbrain-II possesses 4096 pieces of AI processors with 512 server nodes. To efficiently train the language model on such a large cluster, we adopt multiple optimization strategies (Figure 3), reaching a peak performance of 366.8 petaflops with mixed precision.

**Figure 3.**
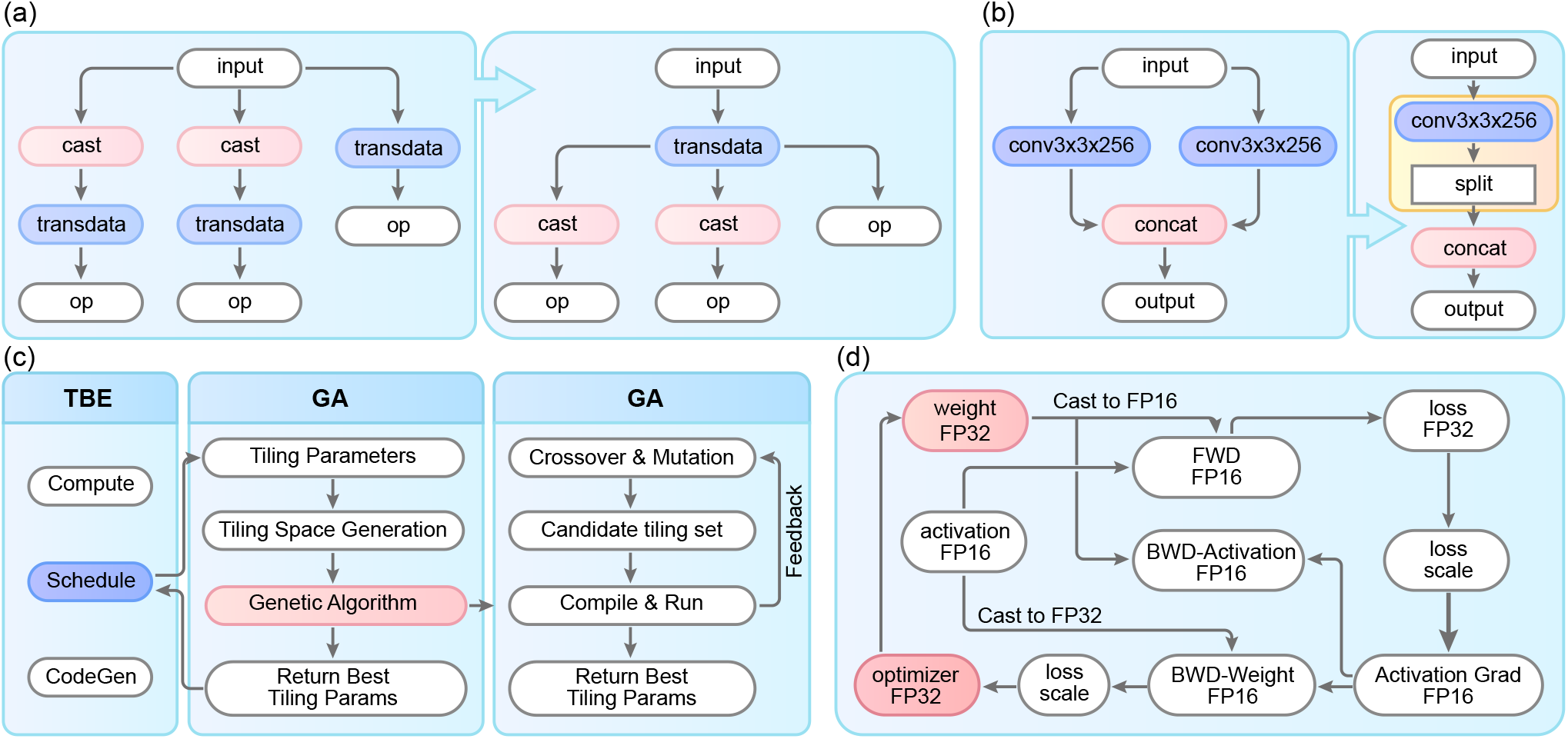
An overview of the employed optimization strategies. (a) Operator fusion. Op means operator. To reduce the redundant memory accesses incurred by the successive execution of many small operators, we integrate multiple transdata operators into one transdata operator. (b) Operator replacement. Conv means convolution. Concat means concatenate. We replace two operators with one simplified operator to reduce the computational cost and model size. (c) Operator auto-tuning. TBE means Tensor Boosting Engine. GA means Genetic Algorithm. We use a genetic algorithm for tuning particular operators by identifying the optimal tiling policies. A well-designed tiling schedule can fully utilize the computing power of the hardware. (d) Mixed precision. All parameters in the model and optimizer are stored in single precision (32-bit), but most of the calculations in this model are performed in half precision (16-bit) to accelerate the training process. This mixed-precision implementation greatly reduces the training latency at the cost of potential overflow due to the limited representation range of half precision.

#### Operator fusion

We run the training task in graph mode and apply pattern-based operator fusion to accelerate the training in this mode. In this work, we perform fusion of the following operators to optimize the ProtFound model: 1) We fuse multiple operators for the forward/backward layer normalization operations and perform calculations on multiple neural processing units (NPU) cores. 2) We fuse the matrix multiplication (*matmul*) operator and the addition (*add*) operator. 3) We fuse the all-reduce operations for all gradients within one Transformer layer into a single operator. These optimizations account for more than 30% of the time consumption.

#### Operator replacement

Operator replacement refers to the replacement of some operators in a model with new operators that are more amenable to online deployment. In this work, we use fast Gaussian Error Linear Unit (GeLU) in place of the original GeLU operator, since the later is not very friendly to NPUs. Such operator replacement can improve the model efficiency by about 10% while maintaining the accuracy performance.

#### Operator auto-tuning

AI computing chips are usually composed of computing units, on-chip storage, data transmission, and other modules. The collaboration among these modules usually significantly affects the computation patterns of operators. The Auto Tune tool of Ascend uses reinforcement learning and genetic algorithm for tuning particular operators by identifying the optimal tiling policies. We use the Auto Tune tool to optimize the *matmul* operator, which accounts for more than 30% of the time consumption.

#### Mixed precision

We further improve the speed performance by using mixed precision schedules. In dozens of layernorm operators, we schedule a reducing sum operation to the Ascend 910 cube core in FP16 and the other remaining operations to the Ascend 910 vector core in FP32 to avoid computation overflow and achieve higher performance. In addition, the embedding and loss calculations are performed in single precision, and the remaining operators are applied in half precision. The optimizer is implemented with single precision. This mixed-precision implementation greatly reduces the training latency at the cost of potential overflow due to the limited representation range of half precision.

### How performance was measured

We perform pretraining of our ProtFound model on Pengcheng Cloudbrain-II with the MindSpore^4^ AI computation framework. We run tests with 8 NPUs per NPU Pod. The tests are scaled from (1 × 8) to (512 × 8) NPUs by powers of 2, and the largest one is assessed on (512 × 8) NPUs at full-scale. Our model reports timings, including epoch times, mini-batch times, and time-to-solution. We measure the full pretraining time-to-solution, scalability, and peak performance at full-scale. We measure the FLOPS for all precisions by using MindInsight, which is a module of MindSpore. We collect floating-point instructions of relevant flavors (that is, addition, multiplication, fused multiply-add, and tensor core operations for FP16, FP32, and FP64) and multiply them by corresponding weighting factors, respectively, to transform them into FLOPS counts. The sum of all these values for all precisions yields our overall mixed-precision FLOPS count. In summary, the criteria used to measure the performance of the ProtFound model are defined as follows:

- **Time-to-solution**, defined as the epoch times of strong scaling.
- **Mini-batch size**, defined as the batch size on a single NPU.
- **Peak performance**, defined as 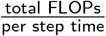.

### Performance results

#### Strong scaling performance

The strong scalability of the pretraining process is measured in terms of the epoch times for 1 to 512 nodes of Pengcheng Cloudbrain-II, as shown in Figure 4. For the strong scaling assessment, the total size of the problem remains the same, i.e., the number of protein sequences used for the ProtFound model pretraining is kept constant at approximately 408 million. The measured strong scaling, shown as a solid line, almost coincides with the optimal strong scaling, shown as a dotted line, which demonstrates that the strong scaling performance is nearly perfect for 1 to 512 nodes. With the performance for 1 node as the baseline, the parallel efficiency at 512 nodes is approximately 96.46%, and the speedup reaches about 493.9×. In addition, the peak performance reaches 366.81 PFLOPS, and the time-to-solution is 9.1 minutes when scaled to 512 nodes in mixed-precision, which enables rapid deployment and iteration of variant generation models.

**Figure 4.**
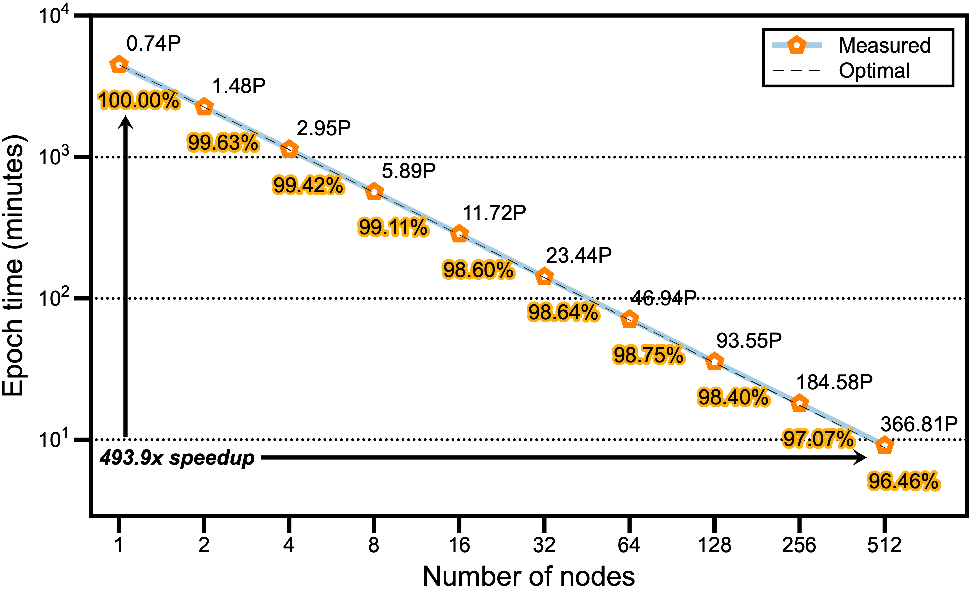
Strong scaling performance of the ProtFound model pretraining for a constant total problem size of approximately 408 million protein sequences. Each data point is labeled with the PFLOPS and parallel efficiency for the corresponding node count. The black dotted line represents the optimal scaling performance for reference.

#### Weak scaling performance

As shown in Figure 5, the weak scaling performance of pretraining the ProtFound model on Pengcheng Cloudbrain-II is also assessed. Unlike the strong scaling case, the problem size per node in the weak scaling test is kept constant at 640 thousand protein sequences. Here, the I/O operations are the saving of checkpoints and trained models. Even if the I/O time is included, the degradation in performance at high node is still slight. Specifically, the parallel efficiency for weak scaling from 1 to 512 nodes slightly reduces from 96.73% to 95.57%, and the utilization also remains stable, reducing from 34.99% to 33.54%. In addition, the peak performance reaches 366.86 PFLOPS (34.99% of Peak) when the I/O time is removed. In summary, for the pretraining of the ProtFound model on Pengcheng Cloudbrain-II, the optimized model scales well to the entire supercomputer.

**Figure 5.**
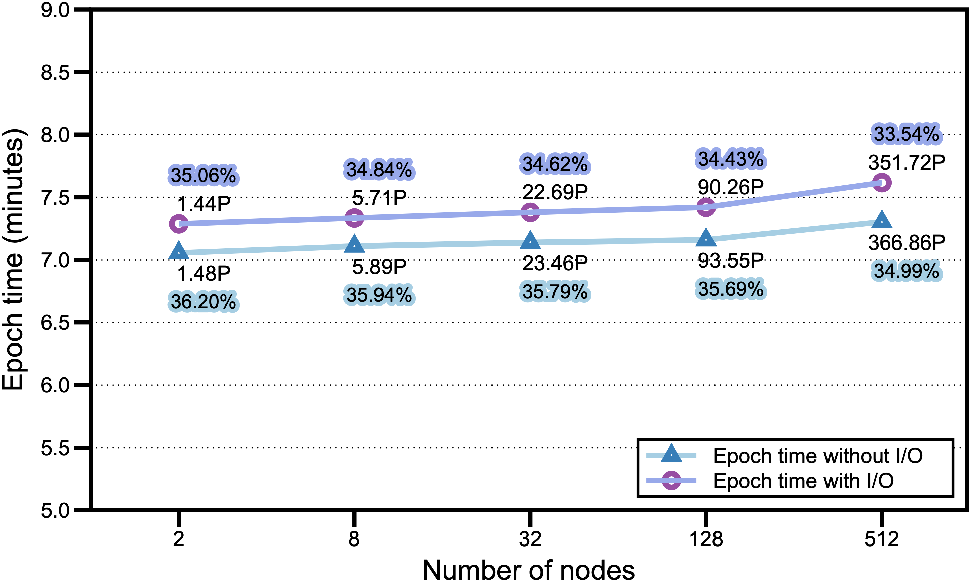
Weak scaling performance of the ProtFound model pretraining for a constant problem size of 640 thousand protein sequences per node. Each data point is labeled with the PFLOPS and utilization for the corresponding node count. Here, the I/O operations include the storage of checkpoints and trained models.

#### In silico validation of RBD mutations of VOCs

The variations of concern (VOCs) that have emerged to date include B.1.1.7 (Alpha), B.1.351 (Beta), P.1 (Gamma), B.1.617.2 (Delta), and B.1.1.529 (Omicron). Omicron, the currently most widespread VOC, exhibits a several-fold accumulation of variants compared with the first four VOCs. Considering the significant difference between the variants before and after the appearance of Omicron, we simulate and verify the RBD mutation process with Omicron as the dividing line as shown in Figure 6.

**Figure 6.**
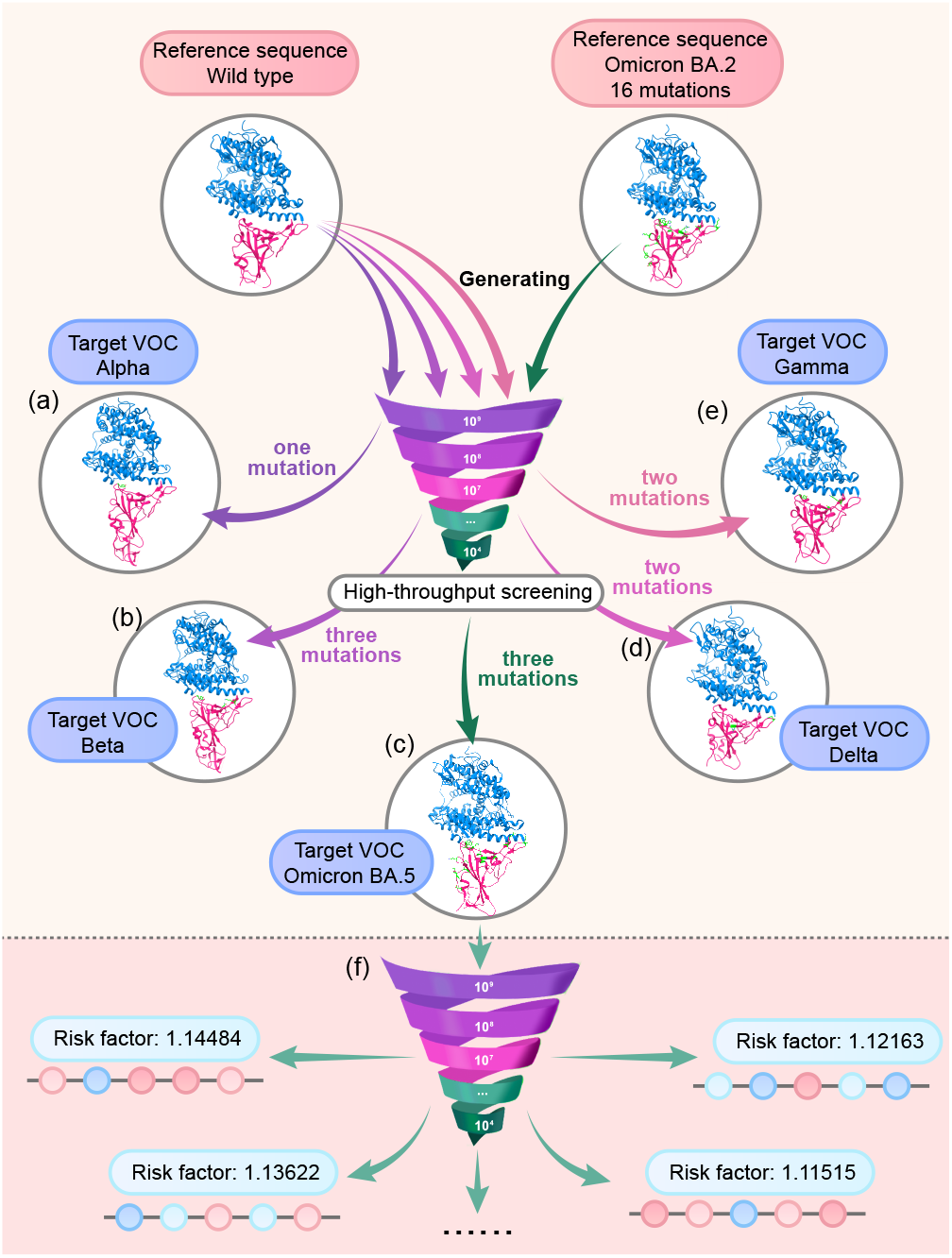
The validation scheme for RBD mutations of VOCs and potential high-risk variants prediction. (a), (b), (c), (d) Four VOCs before Omicron, i.e., Alpha, Beta, Gamma, and Delta, are simulated from wild type to themselves. (e) Omicron BA.5, a latest subvariant of Omicron, is simulated from Omicron BA.2 to itself. (f)Potential high-risk variants prediction. Omicron BA.5 is adopted as the reference sequence. After the high-throughput screening of hACE2 binding and antibody binding, the risk factor is calculated based on mutations relevant to fitness.

For SARS-CoV-2 mutation simulation before Omicron, we validate the predictive ability of MCVP by simulating the mutational changes from the wild type^5^ to the four VOCs (Alpha, Beta, Gamma, and Delta). According to the pathogenic progression of SARS-CoV-2 (Callaway et al. 2022) based on the data from NextStrain^6^, these four VOCs have a parallel evolutionary relationship. Therefore, the starting sequence used to verify the evolutionary route is selected as wild type. The sequences used to fine-tune the model are chosen based on the time when each VOC was first detected. The first detected times and locations of the four VOCs before Omicron are identified via Wikipedia^7^. We segment the data downloaded from GISAID in accordance with the times corresponding to each VOC. For example, Alpha was first reported in September 2020, and we therefore take the data from those submitted before September 2020 as the training sequences for fine-tuning ProtFound to predict the emergence of Alpha. Next we adopt the wild type as the reference sequence for the mutation generation process. After the RBD mutation generation and high-throughput screening, we check the mutated sites to determine if the RBD of Alpha has appeared in the screened RBD mutations. If it appears, the mutation simulation from wild type to Alpha is complete. Otherwise, the filtered RBD mutations are used for iteratively fine-tuning of ProtFound until the RBD of Alpha is generated. Following this simulation method, we have successfully generated the RBDs of the four VOCs (Alpha, Beta, Gamma, and Delta) from the RBD of wild type.

To simulate the evolution of Omicron, we select Omicron BA.2 as the starting point to perform the virus evolving to generate BA.5 in accordance with the pathogenic progression of SARS-CoV-2 (Callaway et al. 2022). In this simulation, the sequences with submission times between BA.2 and BA.5 are selected to fine-tune ProtFound, and BA.2 is used as the reference sequence at the time of generation. Through fine-tuning and identification, BA.5 has been generated successfully by our workflow.

Table 1 shows the proportion of remaining variants after each round of screening. Among the above five VOCs, the variants mutated towards Omicron BA.5 retain more than 80% of the proportion in both the hACE binding and antibody escape screening, which indicates that the Omicron sublineages tend to remain stable binding affinity and have stronger antibody escape capability.

**Table 1.** High throughput screening of various variants

| Variant | 1st screening* | 2st screening** |
| --- | --- | --- |
| Alpha | 39.8% | 2.0% |
| Beta | 13.3% | 51.3% |
| Gamma | 45.2% | 33.8% |
| Delta | 46.7% | 19.1% |
| Omicron BA.5 | 90.4% | 80.2% |

\* Proportion retained in hACE2 binding screening
\*\* Proportion retained in antibody binding screening

#### Potential high-risk mutation prediction

By simulating the mutation of the RBD, we have comprehensively demonstrated that the proposed MCVP can effectively evolve out the RBDs of the known VOCs. However, the real value of MCVP lies in its ability to predict potential future VOCs, thus assisting targeted drug design and vaccine development.

Omicron has been the dominant variant widely spreading around the world. The phenomenon of intra-VOC evolution has been significant due to the sustained transmission of VOCs, which leading to different descendent lineages. In view of this, a variant tracking system, termed “Omicron subvariants under monitoring”, is added to remind us of lineages that need priority attention and monitoring^8^. In this tracking system, BA.5 sublineages (e.g. BF.7, BF.14, BQ.1), BA.2 sublineages (e.g. BA.2.75, BA.2.75.2), and BA.4 sublineage (BA.4.6) need to be focused at present^9^. In order to demonstrate the potential of MCVP to predict future high-risk variants, we simulate the mutational process of BF.7, BF.14, BQ.1, BA.2.75.2, and BA.4.6. As expected, we have successfully simulated these variants that WHO reminds public health authorities around the world to give priority to.

More importantly, as shown in Figure 6f, we take the latest sublineage of Omicron, i.e. BA.5, as the reference sequence, then generate billions of variants in each round and conduct subsequent high-throughput screening. After evaluation of binding affinity and antibody escape capability, we use the screened sequences to fine-tune ProtFound. After several rounds of iterations, we select a number of potential RBD mutations with high risk that maintain a stable binding affinity with hACE2 and a high antibody escape capability. At this stage, to better evaluate potential VOCs, we calculate the relative risk factor based on mutations identified as being associated with fitness of PyR_0_ (Obermeyer et al. 2022). A variant whose risk factor is greater than 0 may have greater risk than wild type, and a variant whose risk factor is less than 0 may have less risk. As a result, billions of variants can be evaluated quickly for the identification of potential high-risk mutations.

### Implications

#### AI models can successfully generate and identify almost all VOCs

In our experiments, using genomic data submitted before the appearance of each VOCs, we successfully generate and identify all VOCs except Omicron. Given the original Omicron spike sequences, we could also generate the Omicron subvariants that are currently the dominant viral variants throughout the world.

During the iterative mutation generation process, the AI models can prioritize mutations based on their predicted binding affinity and antibody escape, two key factors for viral infectivity. Due to their combinatorial nature, it is impossible to experimentally measure the binding affinity changes among all possible RBD mutations (20^201^) and hACE2 or antibodies. Therefore, under the assumption that the deep mutational scanning (DMS) measurements of RBD single mutations might provide reasonable constraints for the RBD to hACE2/antibody binding affinity spaces, we approximate these binding affinity spaces using AI models for prediction of the binding affinities among multiple RBD mutations and hACE2 or antibodies. These AI models are key innovations of the whole workflow.

The fact that our workflow could not generate Omicron despite more than 20 rounds of iteration implies that the mutational features of Omicron are very different from those of other VOCs, since all other VOCs are found after a few rounds of generation.

#### The simulation of SARS-CoV-2 spike mutation is an HPC application

The strategy we used to simulate SARS-CoV-2 spike mutation is dependent on the availability of large-scale genome data (more than 14 million viral genomes as provided by the GISAID database) and a large protein language generation model.

Recent progress in Transformer-based models has enabled the implementation of protein language models capable of generating de novo protein sequences following the principles of natural ones (Ferruz et al. 2022). Inspired by these successes, we pretrain a BERT-like model to learn from millions of viral spike proteins. Our mutation generation workflow heavily relies on the Pengcheng Cloudbrain-II: first, to train the protein language model; second, to iteratively generate new mutations; and third, to evaluate the variants based on AI predictors of: 1) the binding affinity between RBD and hACE2, 2) the antibody escape capability. All the processing steps require an HPC facility, as billions of RBD mutations must be generated in each round and evaluated accordingly.

#### Simulating coronavirus evolution is a new challenge for HPC

The COVID-19 pandemic, caused by SARS-CoV-2, is a stark reminder that coronaviruses remain a major threat to humanity. It is crucial to study the evolution of Coronaviruses to be better prepared for the next pandemic.

SARS-CoV-2 has become the most sequenced virus ever in history, with 14 million SARS-CoV-2 genomes deposited in the GISAID database. The efficiency of simulating these extremely large numbers of closely related genomes to recreate potential histories of past and future virus evolution presents a new challenge for HPC. As proof of concept, in this study, we have initiated the first step toward elucidating the evolution of SARS-CoV-2 VOCs by using only RBD sequences of the SARS-CoV-2 S1 protein. Using all genomes of SARS-CoV-2 in the future, plus other coronavirus genomes, we will be able to perform more reliable simulations to study the evolution of coronaviruses in general and the dynamics of viral transmission across animal species. Meeting the computational requirements of such simulations will require some of the finest HPC systems built to date.

#### SARS-CoV-2 mutation is a serious threat

It has been estimated that an infected person could carry 10^9^ to 10^12^ SARS-CoV-2 virions (Sender et al. 2021). Since the initial outbreak of COVID-19, there have been more than 645 million infections as of December 2022^10^. The potential mutation space for SARS-CoV-2 is thus approximately 6 × 10^17^ to 10^20^. The experimentally deduced spontaneous mutation rate of SARS-CoV-2 is 1.3 × 10^−6^ ± 0.2 × 10^*−*6^ per base per infection cycle (Amicone et al. 2022), which is heterogeneous throughout the genome. Taking all these numbers together, it is not too difficult to conclude that every single base mutation is being generated de-novo and transmitted to a new host every day (Sender et al. 2021). It is therefore extremely important to be able to simulate the viral mutation process and rapidly identify potential VOCs, which is essentially what we have demonstrated in this work through the state-of-the-art AI technology combined with the cutting-edge HPC hardware - the Pengcheng Cloudbrain-II. Any successful prediction of future VOCs of SARS-CoV-2 is not just good scientific research, but can prevent unnecessary deaths.

Further details of this paper will be published later.

## Acknowledgements

We appreciate the useful discussions with Ming Li and Peng Zhou.

## Notes

1. https://covid19.who.int/
2. https://www.who.int/activities/tracking-SARS-CoV-2-variants/
3. https://gisaid.org/
4. https://www.mindspore.cn/en
5. EPI_ISL ID: EPI_ISL_402124
6. https://nextstrain.org/
7. https://en.wikipedia.org/wiki/SARS-CoV-2
8. https://www.who.int/activities/tracking-SARS-CoV-2-variants
9. https://www.cbsnews.com/news/covid-variants-ba46-bf7-ba275-rise-cdc-tracking/
10. https://covid19.who.int/

## References

Amicone M, Borges V, Alves MJ, Isidro J, Zeé-Zeé L, Duarte S, Vieira L, Guiomar R, Gomes JP and Gordo I (2022) Mutation rate of SARS-CoV-2 and emergence of mutators during experimental evolution. Evolution, Medicine, and Public Health 10(1): 142–155.

Beiko RG and Charlebois RL (2007) A simulation test bed for hypotheses of genome evolution. Bioinformatics 23(7): 825–831.

Belouzard S, Chu VC and Whittaker GR (2009) Activation of the SARS coronavirus spike protein via sequential proteolytic cleavage at two distinct sites. Proceedings of the National Academy of Sciences 106(14): 5871–5876.

Callaway E et al. (2022) Are covid surges becoming more predictable? Nature 605(7909): 204–206.

Chen J, Wang R, Wang M and Wei GW (2020) Mutations strengthened SARS-CoV-2 infectivity. Journal of Molecular Biology 432(19): 5212–5226.

Chi X, Yan R, Zhang J, Zhang G, Zhang Y, Hao M, Zhang Z, Fan P, Dong Y, Yang Y et al. (2020) A neutralizing human antibody binds to the n-terminal domain of the spike protein of SARS-CoV-2. Science 369(6504): 650–655.

Coronaviridae Study Group of the International Committee on Taxonomy of Viruses (2020) The species severe acute respiratory syndrome-related coronavirus: classifying 2019-nCoV and naming it SARS-CoV-2. Nature Microbiology 5(4): 536–544.

De Maio N, Boulton W, Weilguny L, Walker CR, Turakhia Y, Corbett-Detig R and Goldman N (2022) phastsim: efficient simulation of sequence evolution for pandemic-scale datasets. PLoS Computational Biology 18(4): e1010056.

Devlin J, Chang MW, Lee K and Toutanova K (2018) Bert: Pre-training of deep bidirectional transformers for language understanding. arXiv preprint arXiv:1810.04805.

Drosten C, Guünther S, Preiser W, Van Der Werf S, Brodt HR, Becker S, Rabenau H, Panning M, Kolesnikova L, Fouchier RA et al. (2003) Identification of a novel coronavirus in patients with severe acute respiratory syndrome. New England Journal of Medicine 348(20): 1967–1976.

Duffy S (2018) Why are RNA virus mutation rates so damn high? PLoS Biology 16(8): e3000003.

Elnaggar A, Heinzinger M, Dallago C, Rehawi G, Wang Y, Jones L, Gibbs T, Feher T, Angerer C, Steinegger M, Bhowmik D and Rost B (2022) ProtTrans: Toward understanding the language of life through self-supervised learning. IEEE Transactions on Pattern Analysis and Machine Intelligence 44(10): 7112–7127.

Ewing G and Hermisson J (2010) MSMS: a coalescent simulation program including recombination, demographic structure and selection at a single locus. Bioinformatics 26(16): 2064–2065.

Ferruz N, Schmidt S and Hoöcker B (2022) ProtGPT2 is a deep unsupervised language model for protein design. Nature Communications 13(1): 1–10.

Fletcher W and Yang Z (2009) Indelible: a flexible simulator of biological sequence evolution. Molecular Biology and Evolution 26(8): 1879–1888.

Gallagher TM and Buchmeier MJ (2001) Coronavirus spike proteins in viral entry and pathogenesis. Virology 279(2): 371–374.

Goyal P, Dollaár P, Girshick R, Noordhuis P, Wesolowski L, Kyrola A, Tulloch A, Jia Y and He K (2017) Accurate, large minibatch SGD: Training imagenet in 1 hour. arXiv preprint arXiv:1706.02677.

He X, Xue F, Ren X and You Y (2021) Large-scale deep learning optimizations: A comprehensive survey. arXiv preprint arXiv:2111.00856.

Hoffer E, Hubara I and Soudry D (2017) Train longer, generalize better: closing the generalization gap in large batch training of neural networks. Advances in neural information processing systems 30.

Hoffmann M, Kleine-Weber H, Schroeder S, Kruüger N, Herrler T, Erichsen S, Schiergens TS, Herrler G, Wu NH, Nitsche A et al. (2020) SARS-CoV-2 cell entry depends on ACE2 and TMPRSS2 and is blocked by a clinically proven protease inhibitor. Cell 181(2): 271–280.

Howard J and Ruder S (2018) Universal language model fine-tuning for text classification. arXiv preprint arXiv:1801.06146.

Hudson RR (2002) Generating samples under a Wright-Fisher neutral model of genetic variation. Bioinformatics 18(2): 337–338.

Ito K, Piantham C and Nishiura H (2021) Predicted dominance of variant Delta of SARS-CoV-2 before Tokyo olympic games, Japan, July 2021. Eurosurveillance 26(27): 2100570.

Keskar NS, Mudigere D, Nocedal J, Smelyanskiy M and Tang PTP (2016) On large-batch training for deep learning: Generalization gap and sharp minima. arXiv preprint arXiv:1609.04836.

Lamers MM and Haagmans BL (2022) SARS-CoV-2 pathogenesis. Nature Reviews Microbiology : 1–15.

Lan Z, Chen M, Goodman S, Gimpel K, Sharma P and Soricut R (2019) ALBERT: A lite BERT for self-supervised learning of language representations. arXiv preprint arXiv:1909.11942.

Laval G and Excoffier L (2004) SIMCOAL 2.0: a program to simulate genomic diversity over large recombining regions in a subdivided population with a complex history. Bioinformatics 20(15): 2485–2487.

Li J, Wu YN, Zhang S, Kang XP and Jiang T (2022) Deep learning based on biologically interpretable genome representation predicts two types of human adaptation of SARS-CoV-2 variants. Briefings in Bioinformatics 23(3): bbac036.

Li W, Moore MJ, Vasilieva N, Sui J, Wong SK, Berne MA, Somasundaran M, Sullivan JL, Luzuriaga K, Greenough TC et al. (2003) Angiotensin-converting enzyme 2 is a functional receptor for the SARS coronavirus. Nature 426(6965): 450–454.

Li Z, Wallace E, Shen S, Lin K, Keutzer K, Klein D and Gonzalez J (2020) Train big, then compress: Rethinking model size for efficient training and inference of transformers. In: International Conference on Machine Learning. pp. 5958–5968.

Liu Y, Ott M, Goyal N, Du J, Joshi M, Chen D, Levy O, Lewis M, Zettlemoyer L and Stoyanov V (2019) RoBERTa: A robustly optimized BERT pretraining approach. arXiv preprint arXiv:1907.11692.

Lu G, Wang Q and Gao GF (2015) Bat-to-human: spike features determining ‘host jump’of coronaviruses SARS-CoV, MERS-CoV, and beyond. Trends in Microbiology 23(8): 468–478.

Lu R, Zhao X, Li J, Niu P, Yang B, Wu H, Wang W, Song H, Huang B, Zhu N et al. (2020) Genomic characterisation and epidemiology of 2019 novel coronavirus: implications for virus origins and receptor binding. The Lancet 395(10224): 565–574.

Mohamed T, Sayed S, Salah A and Houssein EH (2021) Next generation sequence prediction intelligent system for SARS-CoV-2 using deep learning neural network. In: 2021 17th International Computer Engineering Conference (ICENCO). IEEE, pp. 88–93.

Obermeyer F, Jankowiak M, Barkas N, Schaffner SF, Pyle JD, Yurkovetskiy L, Bosso M, Park DJ, Babadi M, MacInnis BL et al. (2022) Analysis of 6.4 million SARS-CoV-2 genomes identifies mutations associated with fitness. Science 376(6599): 1327–1332.

Ofer D, Brandes N and Linial M (2021) The language of proteins: Nlp, machine learning & protein sequences. Computational and Structural Biotechnology Journal 19: 1750–1758.

Osawa K, Tsuji Y, Ueno Y, Naruse A, Foo C and Yokota R (2022) Scalable and practical natural gradient for large-scale deep learning. IEEE Transactions on Pattern Analysis and Machine Intelligence 44(1): 404–415.

Pucci F and Rooman M (2021) Prediction and evolution of the molecular fitness of SARS-CoV-2 variants: Introducing SpikePro. Viruses 13(5): 935.

Rambaut A and Grass NC (1997) Seq-Gen: an application for the Monte Carlo simulation of DNA sequence evolution along phylogenetic trees. Bioinformatics 13(3): 235–238.

Rives A, Meier J, Sercu T, Goyal S, Lin Z, Liu J, Guo D, Ott M, Zitnick CL, Ma J et al. (2021) Biological structure and function emerge from scaling unsupervised learning to 250 million protein sequences. Proceedings of the National Academy of Sciences 118(15): e2016239118.

Sender R, Bar-On YM, Gleizer S, Bernshtein B, Flamholz A, Phillips R and Milo R (2021) The total number and mass of SARS-CoV-2 virions. Proceedings of the National Academy of Sciences 118(25): e2024815118.

Shchur V, Spirin V, Sirotkin D, Burovski E, De Maio N and Corbett-Detig R (2022) Vgsim: scalable viral genealogy simulator for global pandemic. PLOS Computational Biology 18(8): e1010409.

Shu Y and McCauley J (2017) GISAID: Global initiative on sharing all influenza data–from vision to reality. Eurosurveillance 22(13): 30494.

Simmons G, Zmora P, Gierer S, Heurich A and Poöhlmann S (2013) Proteolytic activation of the SARS-coronavirus spike protein: cutting enzymes at the cutting edge of antiviral research. Antiviral Research 100(3): 605–614.

Sipos B, Massingham T, Jordan GE and Goldman N (2011) PhyloSim-Monte Carlo simulation of sequence evolution in the R statistical computing environment. BMC Bioinformatics 12(1): 1–6.

Starr TN, Greaney AJ, Hilton SK, Ellis D, Crawford KH, Dingens AS, Navarro MJ, Bowen JE, Tortorici MA, Walls AC et al. (2020) Deep mutational scanning of SARS-CoV-2 receptor binding domain reveals constraints on folding and ACE2 binding. Cell 182(5): 1295–1310.

Strait BJ and Dewey TG (1996) The Shannon information entropy of protein sequences. Biophysical Journal 71(1): 148–155.

Su S, Wong G, Shi W, Liu J, Lai AC, Zhou J, Liu W, Bi Y and Gao GF (2016) Epidemiology, genetic recombination, and pathogenesis of coronaviruses. Trends in Microbiology 24(6): 490–502.

Suzek BE, Huang H, McGarvey P, Mazumder R and Wu CH (2007) UniRef: comprehensive and non-redundant uniprot reference clusters. Bioinformatics 23(10): 1282–1288.

Trifonov EN (2009) The origin of the genetic code and of the earliest oligopeptides. Research in Microbiology 160(7): 481–486.

Walls AC, Park YJ, Tortorici MA, Wall A, McGuire AT and Veesler D (2020) Structure, function, and antigenicity of the SARS-CoV-2 spike glycoprotein. Cell 181(2): 281–292.

Wan Y, Shang J, Graham R, Baric RS and Li F (2020) Receptor recognition by the novel coronavirus from Wuhan: an analysis based on decade-long structural studies of SARS coronavirus. Journal of Virology 94(7): e00127–20.

Wrapp D, Wang N, Corbett KS, Goldsmith JA, Hsieh CL, Abiona O, Graham BS and McLellan JS (2020) Cryo-EM structure of the 2019-nCoV spike in the prefusion conformation. Science 367(6483): 1260–1263.

Wu C, Liu Y, Yang Y, Zhang P, Zhong W, Wang Y, Wang Q, Xu Y, Li M, Li X et al. (2020) Analysis of therapeutic targets for SARS-CoV-2 and discovery of potential drugs by computational methods. Acta Pharmaceutica Sinica B 10(5): 766–788.

Wu Y, Xu S, Yau ST and Wu Y (2021) Phylotransformer: A discriminative model for mutation prediction based on a multi-head self-attention mechanism. arXiv preprint arXiv:2111.01969.

Yang Z, Dai Z, Yang Y, Carbonell J, Salakhutdinov RR and Le QV (2019) XLNet: Generalized autoregressive pretraining for language understanding. Advances in neural information processing systems 32.

Yin Y and Wunderink RG (2018) MERS, SARS and other coronaviruses as causes of pneumonia. Respirology 23(2): 130–137.

Yu L, Tanwar DK, Penha EDS, Wolf YI, Koonin EV and Basu MK (2019) Grammar of protein domain architectures. Proceedings of the National Academy of Sciences 116(9): 3636–3645.

Zaki AM, Van Boheemen S, Bestebroer TM, Osterhaus AD and Fouchier RA (2012) Isolation of a novel coronavirus from a man with pneumonia in Saudi Arabia. New England Journal of Medicine 367(19): 1814–1820.

Zhou P, Yang XL, Wang XG, Hu B, Zhang L, Zhang W, Si HR, Zhu Y, Li B, Huang CL et al. (2020) A pneumonia outbreak associated with a new coronavirus of probable bat origin. Nature 579(7798): 270–273.

